# NDI: A platform-independent data interface and database for neuroscience physiology and imaging experiments

**DOI:** 10.1101/2020.05.13.093542

**Authors:** Daniel García Murillo, Ora Rogovin, Yixin Zhao, Shufei Chen, Ziqi Wang, Zoey C. Keeley, Daniel I. Shin, Victor M. Suárez Casanova, Yannan Zhu, Lisandro Martin, Olga Papaemmanouil, Stephen D. Van Hooser

## Abstract

Collaboration in neuroscience is impeded by the difficulty of sharing primary data, results, and software across labs. Here we introduce Neuroscience Data Interface (NDI), a platform-independent standard that allows an analyst to use and create software that functions independently from the format of the raw data or the manner in which the data is organized into files. The interface is rooted in a simple vocabulary that describes common apparatus and storage devices used in neuroscience experiments. Results of analyses – and analyses of analyses – are stored as documents in a scalable, queryable database that stores the relationships and history among the experiment elements and documents. The interface allows the development of an application ecosystem where applications can focus on calculation rather than data format or organization. This tool can be used by individual labs to exchange and analyze data, and it can serve to curate neuroscience data for searchable archives.

## Introduction

Despite its importance, collaboration and sharing of data and primary results is very difficult in the neurosciences, particularly for physiology experiments. At present, physiology experiments are usually performed on custom experimental rigs that acquire data in unique, creative, and idiosyncratic ways. Neurophysiology or neuroimaging rigs often employ several pieces of equipment from different eras of time and with vastly different degrees of engineering refinement. Each data acquisition (DAQ) device on a rig usually has its own sampling rate, clock, and means of storing data to disk. On top of this physical heterogeneity are at least 2 types of digital heterogeneity: the digital format of the data, that typically varies from device to device, and the organization of data and metadata into files or folders, that differs greatly from device to device and from lab to lab.

While the current state of affairs allows for significant creativity on the measurement side of experiments, it presents substantial challenges for data analysis and its reproducibility. Most laboratories cannot analyze the data of other laboratories without perhaps a month or more of effort writing conversion software (Teeters et al., 2008; Garcia et al., 2014; Wiener et al., 2016; Rübel et al., 2019; Sprenger et al., 2019). This barrier has meant that most labs or investigators write their own analysis software that they test themselves in only a limited manner. Further, this barrier impedes the development and utility of common, best-of-breed analysis packages that are dedicated to analyzing certain classes of data (Wiener et al., 2016). There are some important efforts to develop file format standards (Teeters et al., 2015; Rübel et al., 2019) that, if followed, would allow for the development of these packages. However, these standards typically require users to first convert their data into the common format, which is itself a barrier to adoption. Heretofore, these packages have been used by relatively few labs, although this situation is improving. It would be ideal to have a tool that allows an analyst to quickly read and analyze data regardless of whether it is organized idiosyncratically or stored in standardized container formats.

Here, we introduce a new approach that allows the development of common analysis tools without requiring a common file format: a Neuroscience Data Interface (**NDI**). The interface provides a standard means of specifying and addressing the data that are collected in neuroscience experiments. At the highest level, the interface provides a vocabulary and conceptual framework for specifying recordings and analyses. At the implementation level, the interface contains an extendible set of open source code and interface standards for reading from a variety of data formats and for specifying the manner in which the experimental data is organized on disk. The interface is platform-and computing language-independent. The interface includes a scalable database for storing results of calculations on the raw data, and user-designed or commercial applications can read and write from the database in order to build complex, layered analyses. These database entries are specified using platform-independent metadata that is human- and machine-readable, and database entries can exist on a user’s computer or in the cloud. NDI is designed to serve analysts who want to be able to quickly read data from a variety of collaborators; if it were widely adopted by the community, it also has the capability to act as a data curation and archive system for neuroscience data.

In this article, we demonstrate the interface in a Matlab prototype. Our purpose here is not to showcase a completed system that works at scale, but is instead to propose a solution to the scientific problem about the level of abstraction that is most useful for wide scale curation and sharing of neuroscience data that allows for the development of common tools. We view this as an important scientific problem at the boundaries of computer science, library science, and neuroscience.

## Results

### Concepts and vocabulary – probes, subjects, elements, DAQ systems, and epochs

Before designing a software interface to experiments, we first sought to codify the elements of an experiment using easy concepts and defined terms, in an effort to take inspiration from the graphical user interfaces developed by Xerox PARC and Apple. We define a **probe** to be any instrument that makes a measurement of or produces a stimulus for a **subject**. *Probes* are part of a broader class of experiment items that we term **elements**, which include concrete physical objects like *probes* but also inferred objects that are not observed directly, such as neurons in an extracellular recording experiment, or abstract quantities, such as simulated data, or a model of the information that an animal has about a stimulus at a given time. Each element must have a *subject*, which can be an experimental subject or an inanimate object like a test resister. We define a **DAQ system** as an instrument or a set of instruments that digitally records the measurements or the stimulus history of a *probe*. These *DAQ system*s record data from *probes* each time the *DAQ system*s are switched into record mode, and we use the term **epoch** to signify each of these recording periods.

The conceptual framework of the interface is applied to a simple experimental situation in **Figure 1**. Here, a *probe* (an extracellular electrode) is used to record activity in the cerebral cortex of a *subject*, a ferret. The *probe* is wired to a *DAQ system* (data acquisition system, DAQ), that is turned on and off 3 times, resulting in 3 *epochs* of sampled probe data that is saved to disk. The probe has been given the name `cortex` and a reference number of `1` in metadata, in this case provided by the user.

**Figure 1.**
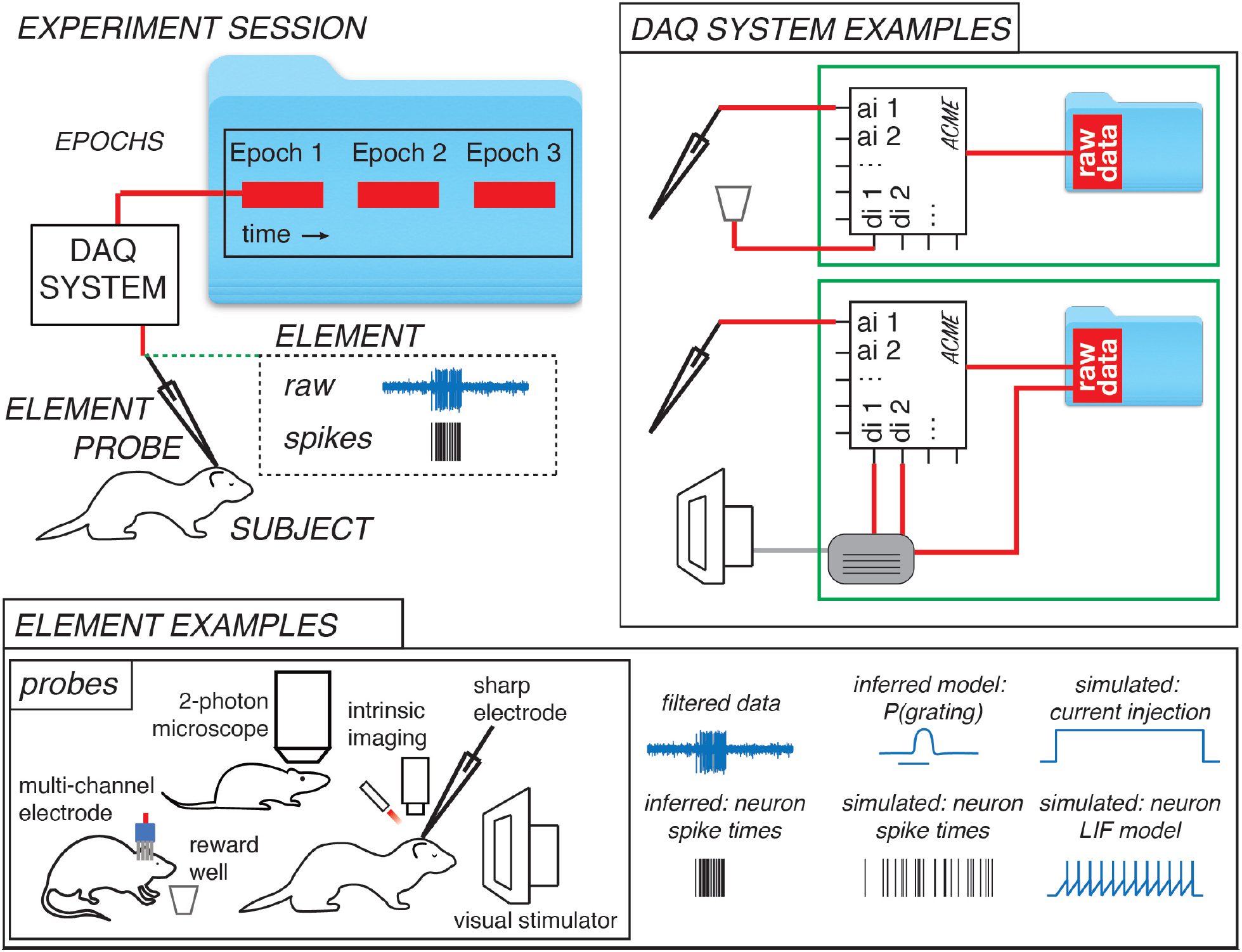
A vocabulary for neuroscience experiments that forms the basis of the Neuroscience Data Interface (NDI). Top-left) An example experiment. A **probe** is any instrument that can make a measurement from or provide stimulation to a **subject**. In this case, an electrode with an amplifier is monitoring signals in cerebral cortex of a ferret and the electrode is a *probe* and the ferret is a *subject*. A **DAQ system** is an instrument that digitally logs the measurements or stimulus history of a *probe*. In this case, a data acquisition system (DAQ) is logging the voltage values produced by the electrode’s amplifier and storing the results in a file on a computer. An **epoch** is an interval of time during which a *DAQ system* is switched on and then off to make a recording. In this case, 3 *epochs* have been sampled. The experiment has additional experiment **elements**. One of these *elements* is a filtered version of the electrode data. A second *element* is a neuron, whose existence and spike times have been inferred by a spike analysis application and recorded in the experiment. **Bottom)** In NDI, a wide variety of experiment items are called *elements*, of which *probes* are a subset. Examples of *probes* include multi-channel extracellular electrodes, reward wells, 2-photon microscopes, intrinsic signal imaging systems, intracellular or extracellular single electrodes, and visual stimulus monitors. Other *elements* include items that are directly linked to *probes*, such as filtered versions of signals, or inferred objects like neurons whose activity are inferred from extracellular recordings or images. Still other *elements* have no physical derivation, such as artificial data or purely simulated data; nevertheless, we want to be able to treat these items identically in analysis pipelines. Finally, *elements* might be the result of complex modeling that depends on many other experiment *elements*, such as an inferred phenomenological model of the amount of information that an animal has about whether a stimulus is a grating. **Top-Right)** *DAQ systems* digitally record *probe* measurements or histories of stimulator activity. In NDI, *DAQ systems* are logical entities, which could correspond physically to a single DAQ device made by a particular company (**top**), or a collection of home-brewed devices that operate together to have the behavior of a single DAQ device (**bottom**). In the bottom example, information from an electrode *probe* and digital triggers from a visual stimulation *probe* are acquired on a single DAQ device, but digital information from both systems (in separate files) is needed to fully describe the activity in each *epoch*.

In this framework, a large variety of experimental apparatus are considered *probes*. Examples of *probes* that make measurements include a whole cell pipette, a sharp electrode, a single channel extracellular electrode, multichannel electrodes with either known or unknown geometries, cameras, 2-photon microscopes, fMRI machines, nose-poke detectors, EMG electrodes, and EEG electrodes. Examples of *probes* that provide stimulation are odor ports, valve-driven interaural cannulae, food reward dispensers, visual stimulus monitors, audio speakers, and stimulating electrodes.

In an experiment, we also deal with items that we do not observe directly, or abstract items, or simulated data. We term all of these items as experiment *elements* (avoiding the term “object” to minimize confusion with the software objects in the implementation). An example of an inferred *element* is the activity of a neuron derived from an extracellular recording. We do not observe the neuron directly, so while we have some certainty that it corresponds to a physical entity, this is really an inference, and different analysts may disagree as to whether it exists. Another type of quantity that we may wish to use in our analysis is a model, such as a calculation of the information that the animal has about a stimulus at a given time. Moreover, we may wish to generate artificial data or simulated data that will go through the same pipelines as experimental data. Thus, experiment *elements* encompass a broad class of items, including *probes*.

To read the data generated by a *probe*, **NDI** must access data from the data acquisition device or devices that recorded the probe, which we term a *DAQ system*. A *DAQ system* can either be a single data acquisition system, such as a data acquisition device made by a major company, or it can describe the collective recordings of a set of these systems, such as a home-brew system that might use a few data acquisition devices at a time. In our own lab, our visual stimulation system relies on data from 2 data acquisition systems (our stimulus computer and a multifunction data acquisition system that records digital triggers), but logically these are treated together as a single *DAQ system* in NDI (**Figure 1**).

Each time a *DAQ system* is switched on and off, an *epoch* of data is logged. The *epochs* are numbered (1, 2, etc) and assigned a unique identifier that never changes, so that the *epoch* can be unambiguously referenced even if other *epochs* are added or deleted later. It is also necessary to specify, for each *epoch*, the mapping between any *probes* that are present and the channels of the *DAQ system* that correspond to the *probes*. Commonly, this information must be specified manually using a data type that we have created, but some multifunction data acquisition systems (such as SpikeGadgets MFDAQs) and file formats include this *epoch* metadata in their native file formats, and this metadata can be processed from the files directly.

With a vocabulary to describe the real-world items in an experimental session, we can describe the necessary computational features of the interface (**Figure 2**). While the specification of the *probes, subjects, elements, DAQ systems*, and *epochs* is sufficient to allow the interface to read the data from the *probes* in the experiment, it would be useful to the analyst and his/her collaborators to have a space to store the results of analyses of this data. This space is provided by the **database** (**Figure 2**), which allows the user to store any type of text or binary data related to the experiment in entries called **documents**. For example, one may have a *document* that stores the responses of a neuron to a family of stimuli, and another *document* that stores the results of a model fit of that neuron’s responses to the stimulus family. Still another *document* might store the aggregate statistics of the responses to all the neurons in a given study. *Documents* in NDI have a human-readable portion and the option of a binary blob, so that they can be understood easily by humans and programs.

**Figure 2.**
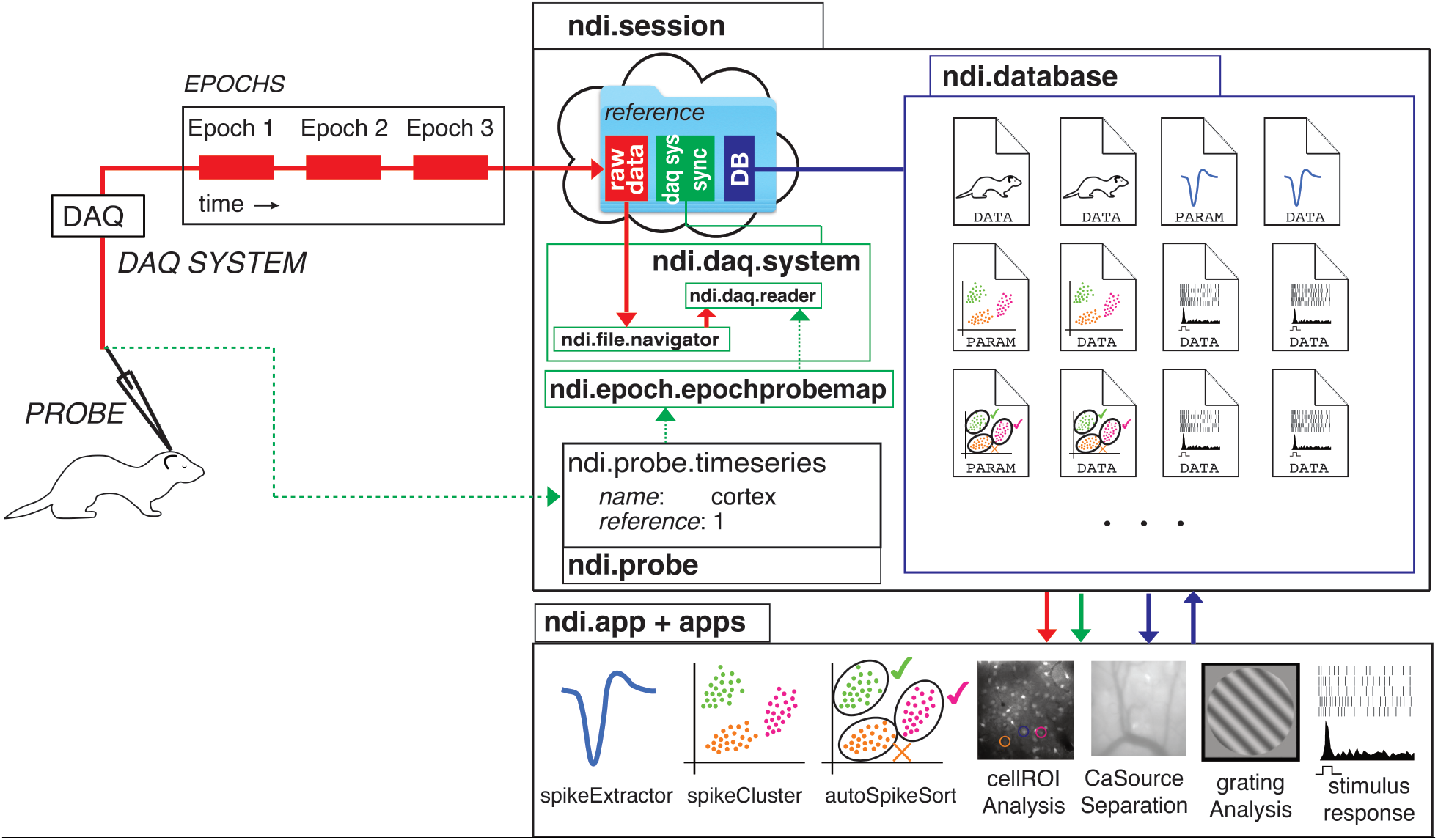
An overview of the Neuroscience Data Interface (NDI). Top-left) The physical experiment from Figure 1. A *probe* (electrode) is used to sample data from the visual cortex of a *subject* ferret. A *DAQ system* digitally logs the measurements. 3 *epochs* of data have been recorded by the *DAQ system*. **Top-right)** An experiment *session* is contained in a software object that has a link to the **raw data** (red), an internal set of NDI objects that have information about *DAQ systems* and synchronization methods (green), and link to a **database** (dark blue). Upon creation, each **ndi.daq.system** object is provided with an **ndi.file.navigator** object, which is a parameterized set of instructions for locating the raw files or links that contain the data for a given *epoch*. Therefore, the same *ndi*.*daq*.*system* can manage data that is organized into *epochs* on disk according to different schemas. Metadata associated with each *epoch*, in a type called **ndi.epoch.epochprobemap**, specifies the *probes* that are present in each recorded *epoch* and indicates the *probe*’s name, a unique reference, and the channel mapping between the *ndi*.*daq*.*system* and the *probe*. This data can be added manually by the user or analyst, or can be read from the *epoch* data files if the *ndi*.*daq*.*system*’s data format or a Laboratory Information Management System (LIMS) encodes this information. The database stores documents, which are platform-independent representations of analyses, analyses of analyses, and NDI internal objects. **Bottom-right)** Applications can use NDI to read raw data and read the results of previous analyses from the *database* and write the results of new analyses back to the *database* as *documents*. The *database* and *documents* therefore support the construction of pipelines that may be linear or integrated. Applications are free to focus on single analysis problems instead of the raw data format or organization of their input.

The *interface* with the *database* allows the creation of an **application ecosystem** (**Figure 2**) that can read the raw data and read and write to the database. For example, one common set of early analyses that must be performed by physiologists examining extracellular data is to identify spike waveforms from the raw data and to make an inference as to which spike waveforms arise from the same neuron(s). The NDI *document schema* specifies a *document* type that includes common spike detection parameters, including threshold algorithm, filter frequencies, the amount of time around each spike to extract, refractory period, etc. These parameters can be used by a variety of spike extraction applications, including the example “*spikeExtractor*” app shown in **Figure 2** but also other related applications that may be developed. There is also a *document schema* for storing extracted spike waveforms and the spike times, and another *schema* for spike shape features. These *documents* can be used by spike sorting applications, such as the example “*spikeCluster*”, to produce assignments of spikes to clusters. One can imagine another application that automatically performs neuron assignment from these clusters (“auto*SpikeSort*”), and so on. The *document schemas* are flexible and expandable, but must contain certain fields that applications can count on being present. In this way, developers and scientists can write applications that perform a particular job well, and mix and match their desired applications. The *database* and *document* schema allows for powerful collaboration across applications, and allows for a healthy competition and interchangeability among applications that perform similar jobs.

The *database* is also designed to allow for the curation and examination of neuroscience data and computations at scale. Because each *database document* contains the identifier of the experimental session, the *documents* can be combined and searched across the cloud so that data and analyses from multiple experiments can be queried, allowing third parties to easily perform analyses or meta analyses of a wide variety of experimental data.

The interface is also meant to be used in a similar manner during on-line evaluation of data and off-line evaluation of data. The data is addressed in the same manner regardless of whether it has been acquired in the last few seconds or a long time ago. This design choice has the advantage that all applications can be used on-line or off-line, and removes the necessity of any second “curation” step before making data available to the world on a data archive. The data can be curated live, during the experiment.

### Implementation -high level

The Neuroscience Data Interface is both an idea, as described above, and an evolving open-source software product that implements the concepts. The current software implementation of NDI has two layers: a high-level layer of core objects that are described here, and a low-level of objects that implement the details of the high-level objects. The separation between the high-level and low-level objects has been made so that the external interface of NDI can be stable, while the open-source products that implement file reading or the database can be switched in and out over time without greatly impacting the user/analyst’s use of the interface. The high-level interface is intended as a sort of “neural data operating system” on which GUIs and other programs can build, but the core of NDI does not define any particular graphical user interface or stipulate the use of any particular underlying database product.

The goal of this paper is to describe the high-level objects in brief so that the ideas of the interface can be discussed or criticized. This paper is not meant to serve as a software tutorial. For tutorials on using the software with neuroscience data, please see the repository of our current software at http://github.com/VH-Lab/NDI-matlab. As of this writing, the Matlab prototype is mature and a Python prototype is still under development.

### Reading from data acquisition systems: ndi.daq.system

An `ndi.daq.system` object is a means of addressing and reading the files that are stored by the DAQ devices that comprise a *DAQ system*. Different high-level subclasses of `ndi.daq.system` allow the user to read from multifunction data acquisition systems (with analog and/or digital channels and sampling rates: `ndi.daq.system_mfdaq`), from imaging systems (with image channels and frames: `ndi.daq.system.image`), or from stimulus systems (with events and parameters: `ndi.daq.system.stimulus`).

All `ndi.daq.system` objects rely on 2 key software objects that determine the `ndi.daq.system` object’s input and output. The first of these is an `ndi.file.navigator` object, which allows the user to specify, with a few parameters, how the system should search for the files that correspond to each recording *epoch*. **Figure 3** shows how different parameters and subclasses of the `ndi.file.navigator` class can be used to navigate the different file organization schemas of different labs. Once the files are found, another software object, the `ndi.daq.reader`, provides the services for reading data from the particular file formats that comprise the epochs.

**Figure 3.**
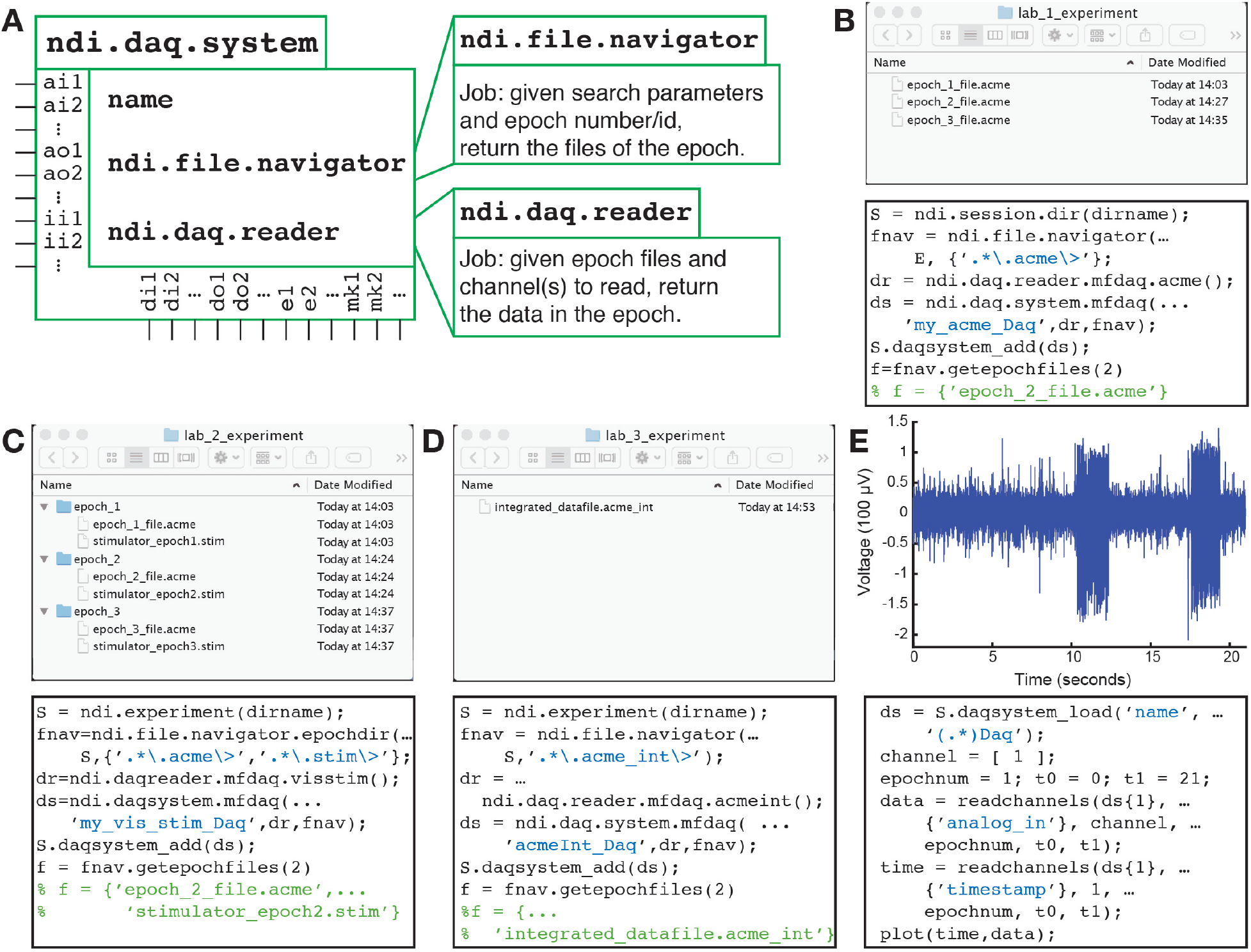
*DAQ systems* allow an analyst to read data in a variety of formats and with a variety of file organizations on disk or in the cloud. All labs begin by initializing the main data management object, an `ndi.session`. **A)** In lab 1, data from an ACME DAQ device (`.acme` files) is organized in a single, flat directory. With a search parameter (the regular expression `.*\.acme\>`), an `ndi.file.navigator` object is instructed how to find the data for each epoch. The file for epoch 2 is requested and shown. **B)** In lab 2, data from a home-brewed configuration using an ACME DAQ device that writes `.acme` files and a custom stimulation system that writes `.stim` files are organized in a single DAQ system. In this lab, data from individual epochs are contained in subdirectories. A subclass `ndi.file.navigator.epochdir` is used to restrict epochs to the contents of subdirectories, and the search parameters indicate that an epoch must have both a `.acme` file and `.stim` file to be valid. **C)** Lab 3 uses an integrated file format, such as that from SpikeGadgets. **D)** After setting up the *DAQ systems*, data for all the labs is read using the same code, which is independent of the file format or the organization on the disk or server.

### Reading from probes: ndi.element and ndi.probe

When an analyst thinks of a *probe* such as an electrode, he or she might think of the *probe* as having the properties of the data acquisition system that records it. For example, we may want to talk about the channels of the electrode, and even casually speak of the “sampling rate” of an electrode despite the fact that it is the *DAQ system* that directly has a sampling rate, not the electrode. The `ndi.element` class, of which `ndi.probe` is a member, allows one to address the *probe* or *element* directly, without regard to the *DAQ system* that acquired it, which is handled behind the scenes by NDI. In order to define a *probe*, it is necessary to functionally define, for each recording *epoch*, a map between the channels of the `ndi.daq.system` and the `ndi.probe` object. This can be done manually with the class `ndi.epoch.epochprobemap`, or can be specified in the data files directly if the *DAQ system* allows it. As shown in **Figure 4**, *probes* can be read by analysis programs without any direct concern about the underlying *DAQ systems* that were employed.

**Figure 4.**
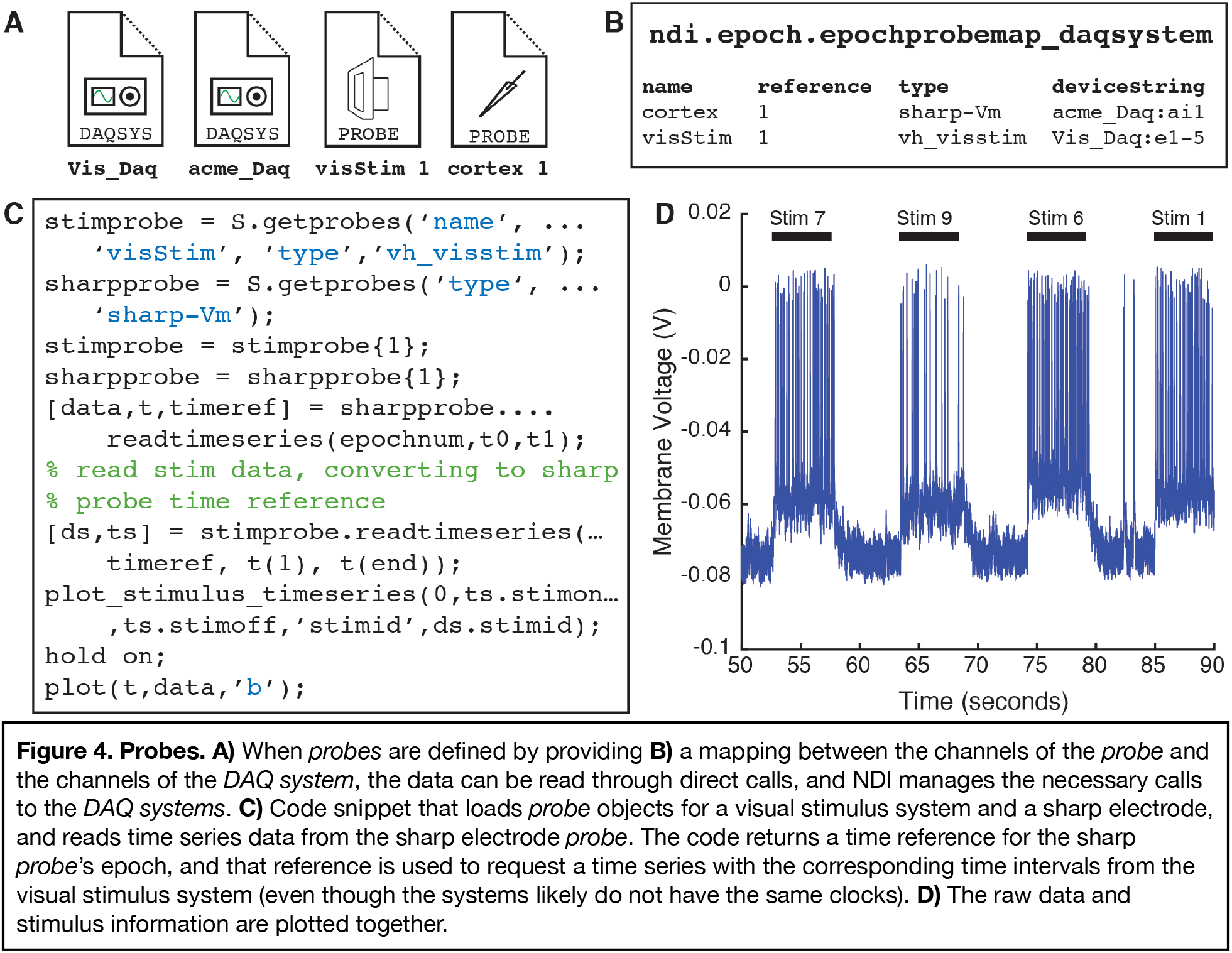
Probes. **A)** When *probes* are defined by providing **B)** a mapping between the channels of the *probe* and the channels of the *DAQ system*, the data can be read through direct calls, and NDI manages the necessary calls to the *DAQ systems*. **C)** Code snippet that loads *probe* objects for a visual stimulus system and a sharp electrode, and reads time series data from the sharp electrode *probe*. The code returns a time reference for the sharp *probe*’s epoch, and that reference is used to request a time series with the corresponding time intervals from the visual stimulus system (even though the systems likely do not have the same clocks). **D)** The raw data and stimulus information are plotted together.

The `ndi.element` class allows many types of data to be treated similarly by software programs. For example, all time series in NDI are members of a subclass called `ndi.element.timeseries`, which can include artificial (test) data, modeled data, filtered data, and so on. In **Figure 5**, the user has created 2 `ndi.element.timeseries` objects from a recording from a sharp electrode: 1 of these *elements* represents the membrane voltage where the spikes have been removed by a median filter, and the other represents the the spiking activity of the cell that is recorded by the sharp electrode. These `ndi.element.timeseries` objects can be passed along to an analysis application (here, our built-in applications `ndi.app.tuning_response` and `ndi.app.oridirtuning`). The epochs of both of these *element* objects are linked back to *epochs* in the *probe*, which are in turn linked to the *epochs* of the *DAQ system*, so that time relationships between other systems, such as the visual stimulus system, are automatically understood for all of the *element* objects derived from *probes*.

**Figure 5.**
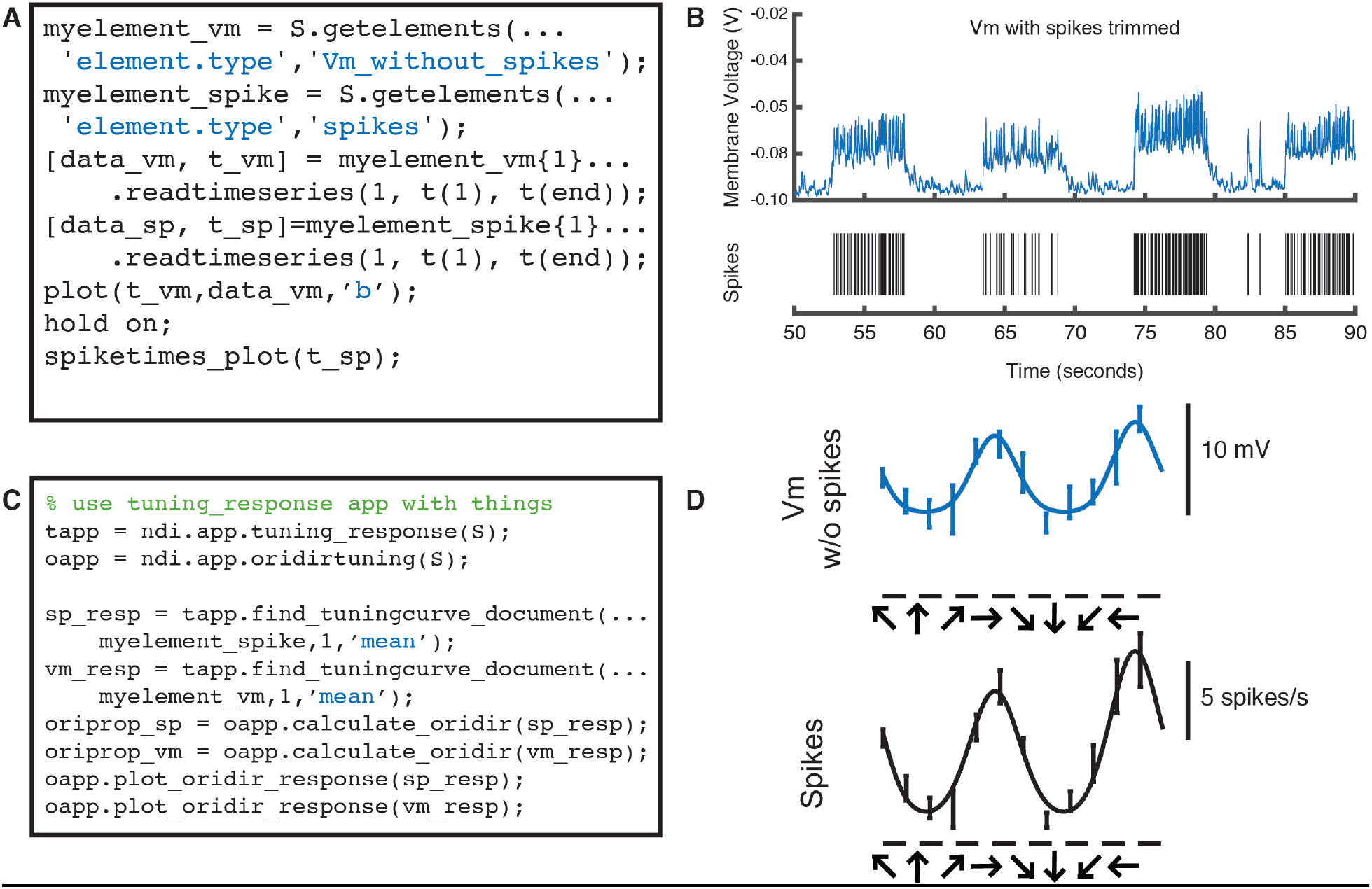
ndi.element objects allow different types of data to go through identical analysis pipelines. **A)** Code that reads and **B)** plots time series data from 2 `ndi.element` objects derived from a single sharp electrode probe: voltage membrane data where spikes have been “chopped” out with a median filter (top) and thresholded spike data (bottom). **C)** The objects can be sent through analysis applications identically and the same type of summary data generated and plotted. **D)** Orientation and direction tuning curves for the subthreshold membrane voltage and spiking activity of the same cell. Note that filtered data, modeled data, or artificial test data can be sent through the same analysis pipelines with `ndi.element`.

### Clocks and time: ndi.time.clocktype,ndi.time.timereference,ndi.time.syncgraph, ndi.time.syncrule

One of the biggest challenges in experiments that involve multiple *DAQ systems* is synchronizing time across devices that have different clocks. In general, data acquisition devices do not share the same clocks: the current time reported by each device will differ from others at any given time, and the drift rate of these clocks differs very slightly in a matter that may alter the timing of samples in long recordings. Many current data standardization schemas sidestep this issue and simply insist that the user must convert all times into a standard clock, and NDI is rare in building clocks and synchronization into the interface.

NDI defines several types of clocks (`ndi.time.clocktype`). The most common type of clock is “device local time” (`dev_local_time`), which means that a *DAQ system* has a local clock that, for each *epoch*, starts a time *t*_*0*_ and ends at a time *t*_*1*_. In most cases, *t*_*0*_ is 0, and *t*_*1*_ is the duration of the recording. Some devices may further keep a “device global” time, so that the device has a sub-millisecond record of the relationship between the *t*_*0*_ of a given recording *epoch* and the *t*_*0*_ of a second recording *epoch* on the same device, but this is unusual. We also define the possibility that a device has a record of some “global experimental time” or that it keeps “universal controlled time” (UTC).

As analysts, we’d like to be able to refer unambiguously to a time *t* on the clock of a given *DAQ* system, and effortlessly know the corresponding time *t’* on the clock of another *DAQ* system. Therefore, built into every call to the function `readtimeseries`, which reads data from a time *t*_*i*_ to a time *t*_*j*_ from an `ndi.element`, `ndi.probe`, or `ndi.daqsystem`, is an input that specifies the time reference (`ndi.time.timereference`) being used. `ndi.time.timereference` objects include the referent (the `ndi.element`, `ndi.probe`, or `ndi.daqsystem` being referred to), the clock type, an *epoch* id (if the `ndi.clocktype` is `dev_local_time`, which is most common), and an offset time.

The system is illustrated in **Figure 4**. Here, the user reads samples from a sharp electrode *probe* using `readtimeseries`, which returns the *time reference* that was used. Next, the user wants to extract stimulus times from the visual stimulus *probe*, which has a different clock. The user simply passes the *time reference* object that was returned from the sharp electrode *probe* to the `readtimeseries` call to the visual stimulus *probe*, and NDI converts the input and output times appropriately so that the output returned is relative to the sharp electrode *probe*’s clock.

The interface solves these conversions from a given clock to another clock by computing paths through a directed graph that contains all recorded *epochs* as nodes and the mappings between *epochs* as edges. The object that performs this computation is called `ndi.time.syncgraph`. The mappings across *epochs* recorded on different *DAQ systems* are typically calculated by examining recordings of the same signal (such as a set of digital triggers) on both *DAQ systems*. One can also specify rules of synchronization (`ndi.time.syncrule`) among devices, and `ndi.time.syncgraph` will automatically calculate possible mappings from its set of `ndi.time.syncrule` objects and solve the paths through the graph. An `ndi.time.syncrule` might specify the channels of 2 *DAQ systems* that record digital triggers in common, or might specify that 2 *DAQ systems* have the same clock if one of their data files is shared between the 2 systems (such that the same DAQ hardware is being used in service of 2 *DAQ systems*). Sometimes, if DAQ systems were not used simultaneously, or if there is no `ndi.time.syncrule`, there is no known mapping between different *epochs*. For example, if a DAQ system only has a local clock, then we usually do not understand the time relationship between subsequent epochs of that system (and usually there is no need to understand this relationship). Example cases of synchronization relationships are shown in **Figure 6** and **Figure 7**.

**Figure 6.**
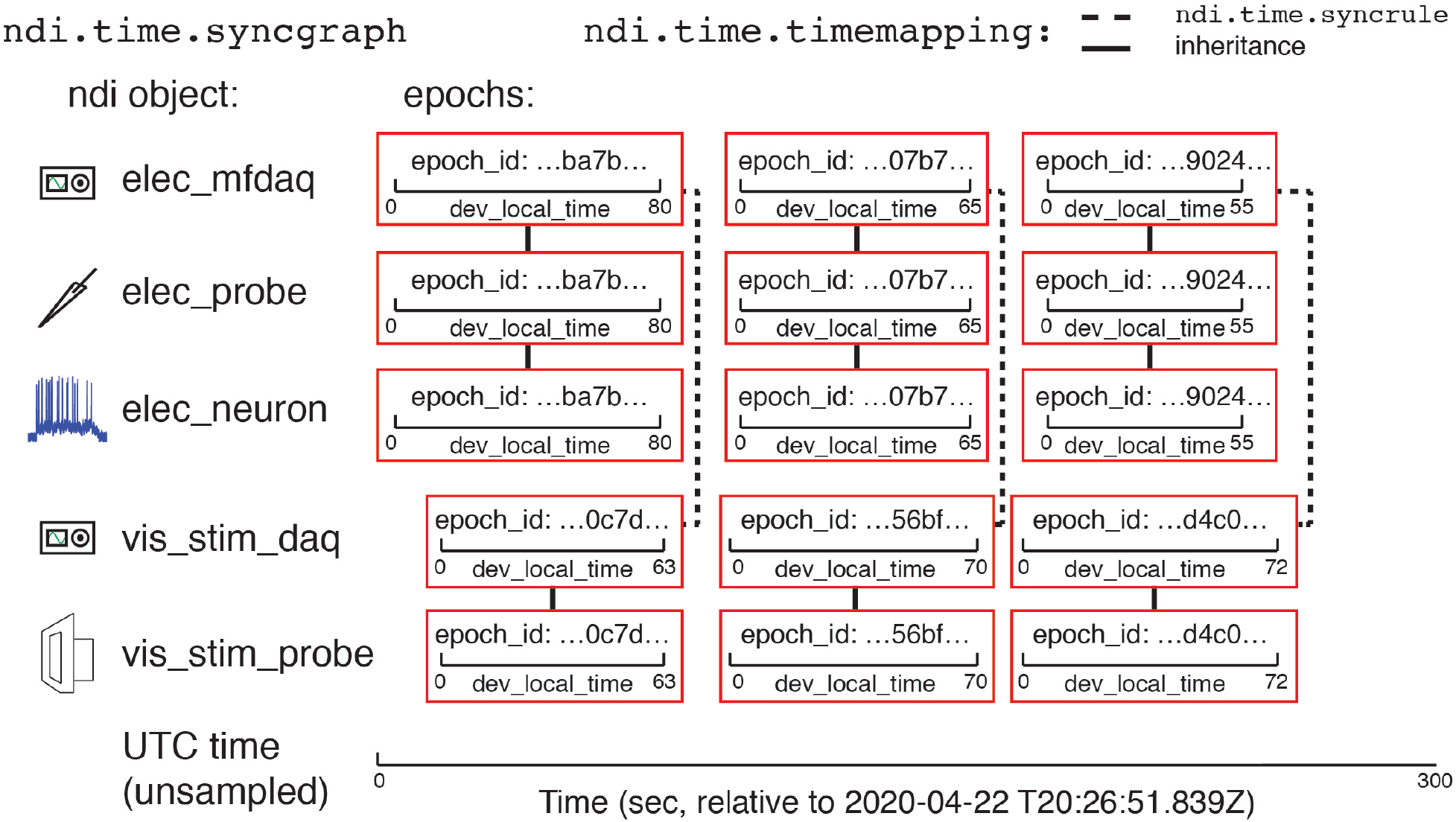
Epochs and ndi.time.syncgraph. Illustration of an example experiment with 2 `ndi.daq.system` objects (`elec_mfdaq` and `vis_stim_daq`) that are each connected to a probe (`elec_probe` and `vis_stim_probe`, respectively). The DAQ systems have their own clocks that are not linked to any global time system. 3 epochs have been recorded by each *DAQ system*. The electrode probe has been analyzed and an `ndi.element` object (a neuron, `elec_neuron`) has been created from it. The clock and time of each of the epochs for the neuron is inherited from its underlying *probe*, which is in turn inherited from the underlying *DAQ system*. The 2 *DAQ systems* each record the same set of digital triggers, and `ndi.time.syncgraph` has used its list of `ndi.time.syncrule` objects to compute a mapping (`ndi.time.timemapping`) between epochs of those DAQ systems. Time can be converted between epochs that are recorded simultaneously on the 2 *DAQ systems*, but we do not know how the other epochs are related to each other, or how any epoch is related to a global time system like universal controlled time (UTC), shown below.

**Figure 7.**
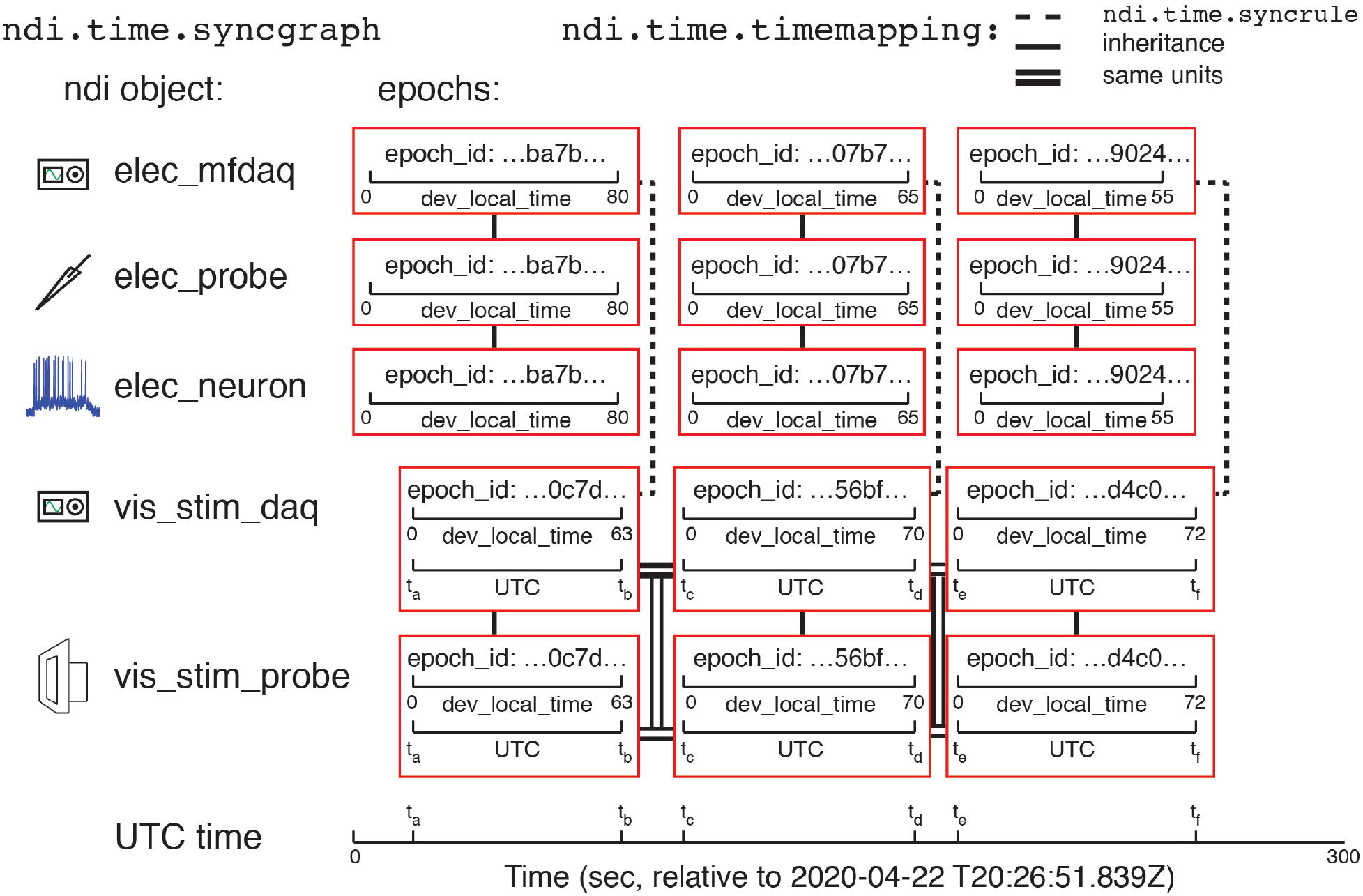
Epochs and ndi.time.syncgraph. Illustration of an example experiment similar to that in **Figure 6**, except that the `vis_stim_daq` *DAQ system* also keeps UTC time in addition to its own local clock. Here, time can be converted among any epoch because there is a mapping between the epochs of `vis_stim_daq` and UTC, and there are `ndi.time.timemapping` mappings between the DAQ system. The time in any epoch can be computed according to the clock of any other epoch, by solving the transformations in the *syncgraph*. The mappings shown are `ndi.time.timemapping` objects built by a) an `ndi.time.syncrule`, b) inheritance (e.g., a *probe* inherits the *epoch* information of the *DAQ system* that acquired it); and c) same units (UTC is a global time system).

### Database, documents: ndi.database and ndi.document

All of the interface that we have described so far is necessary for reading raw electrophysiology or imaging files, but does not allow the user to store the results of analysis in a convenient and well-documented manner. For this purpose, each experiment is linked to a *database* that can be running on the local computer or in the cloud. The database class `ndi.database` provides standardized methods for adding *documents* to the database that conform to a validated, open schema, searching the *database*, and removing *documents* from the *database*.

The fundamental unit of the *database* is the *document*, which is implemented by the software class `ndi.document`. All *documents* include a core structure of fields that describe the unique identifier of the experiment session, the unique identifier of the *document*, the time of creation, the schema of the *document*, and a history of how the *document* was created so that the calculation can be traced back to the raw data or antecedent computations in other *documents. Document schemas* are specified in a platform independent, human-readable format so they can be read and interpreted on any platform and be read and understood by human readers easily. *Document* classes can be composed so that one can build *documents* that refer to common elements (such as *epoch* ids or app properties) in a consistent manner across *documents* (**Figure 8**). Dependencies among *documents* can also be expressed so that relationships among *documents* in a pipeline are clear. Finally, each *document* has its own binary stream that can be used to store large binary data.

**Figure 8.**
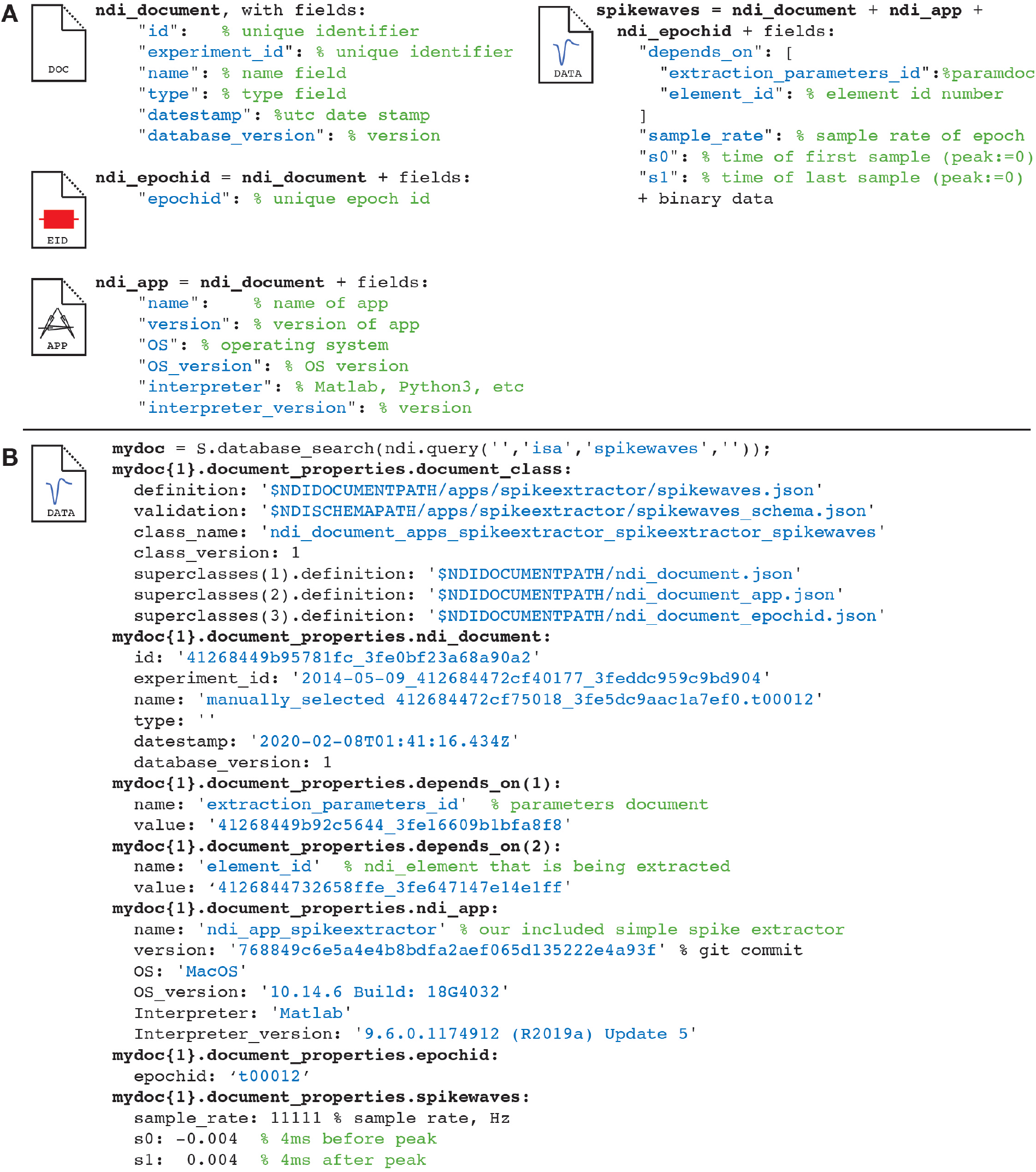
Illustration of ndi_documents and the creation of new classes of ndi_documents by composition. **Top panel)** *Document* definitions, with fields. Several *document* classes are created by composition: for example, the `spikewaves` type has its own fields plus those of document classes `ndi_document`, `ndi_epochid`, and `ndi_app`. **Bottom panel)** A specific spikewaves *document* from a *database*. The *document* includes a description of the *document* definition, a unique ID and timestamp, the app that created it, the parameters that were used, a link to the `ndi.element` that was analyzed and other parameters.

Note that the idea for an extendible, local-or cloud-based database of this type is not new. For example, the open-source program DataJoint (Yatsenko et al., 2015) uses a similar design, although the underlying data are organized into smaller units called tables rather than *documents*. The tables in DataJoint are similar to the substructures of NDI *documents*.

### Analysis pipelines: ndi.app and ndi.query

To understand the power of the interface and the potential app ecosystem, it is useful to examine a simple analysis pipeline. In this pipeline, we will use a simple spike detection app that is included in the base distribution of NDI called `ndi.app.spikeextractor` to detect spikes in sharp electrode data, and then user code to plot the spike shapes.

The steps of the code that produces the pipeline are illustrated in **Figure 9**, along with the *database documents* that are produced at each step. First, the user opens an experiment session and identifies the sharp electrode data for each epoch. The data here has been normalized by subtraction so that the voltage activity during the preceding interstimulus interval (blank screen) is 0. Then, the user creates an instance of the application `ndi.app.spikeextractor` (Step 1), builds a *document* that has a set of parameters that the app will use in identifying spikes, and adds this *document* to the *database* (Step 2). Next, the user calls the app’s `extract` method to find and extract the spike data from the *element*; the results of the extraction, including spike times and spike shapes for each epoch, are added to the *database* as a *document* (Step 3).

**Figure 9.**
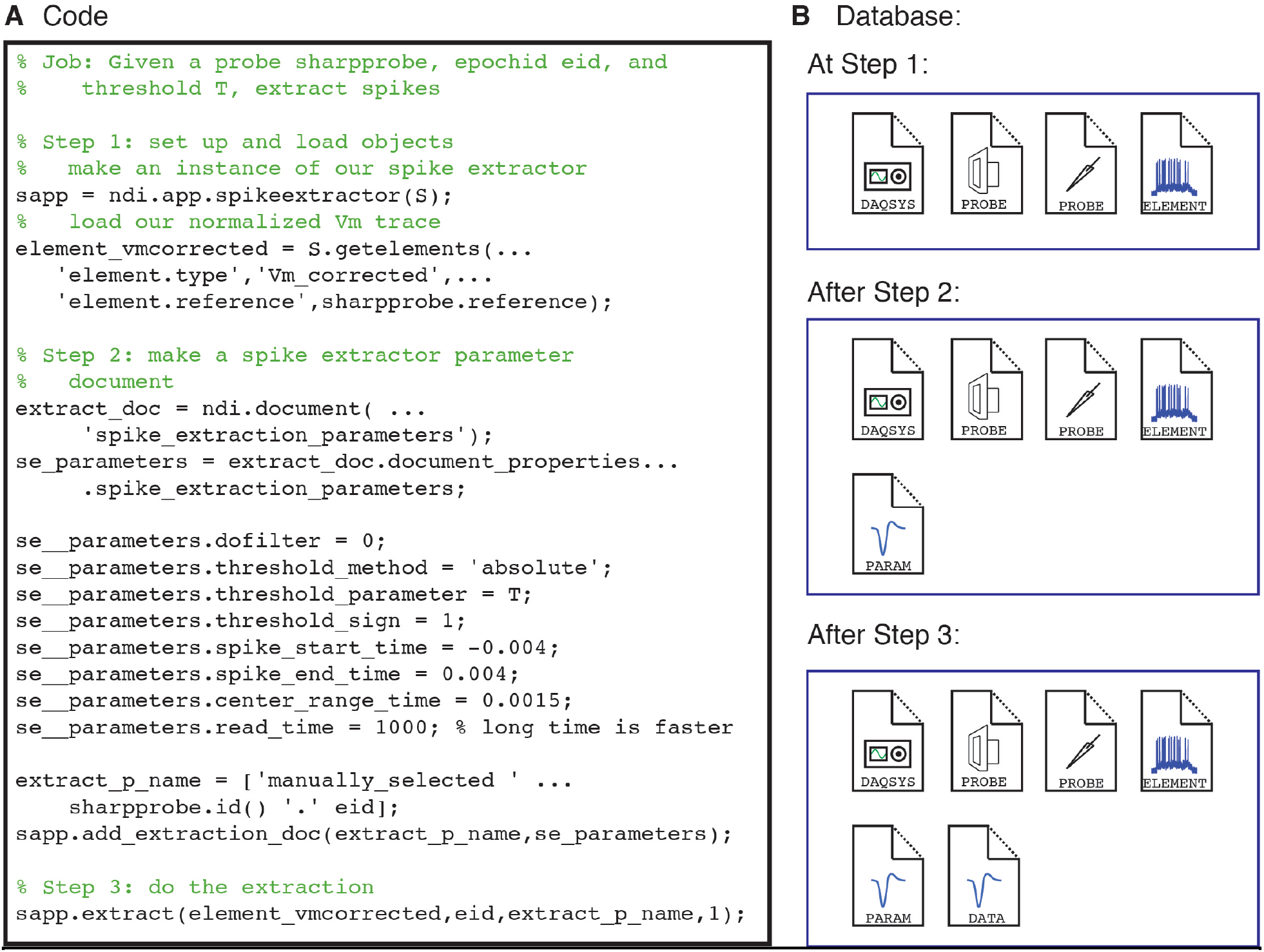
Analysis pipelines build *database documents*. **A)** Code snippet that creates an instance of the NDI spike extractor app (Step 1), creates a *document* that contains the parameters to be used for spike waveform extraction (Step 2), and extracts the spikes (Step 3). **B)** The *database documents* that are present at each Step. Initially, the experiment has an `ndi.daq.system`, 2 *probes* (a visual stimulus system and a sharp electrode), and an `ndi.element` that is a normalized version of the spiking activity. At Step 2, a *document* describing the parameters to be used for spike waveform extraction is added. At Step 3, a *document* describing the extracted spikes is added.

To see what results have been computed, it is necessary to search the *database* for the analysis *documents* that currently exist. The *database documents* can be queried with a search object called `ndi.query`, which allows the user to perform many types of searches. For example, the user can search any text field for several types of matches (exact, partial, regular expression match) or search any number field for several types of matches (equal to, greater than, less than, etc). The user can also search for *documents* of specific types, membership in a particular *session*, and search for *documents* that “depend on” specific other *documents*. **Figure 10** shows a short example of the user using `ndi.query` to check for the existence of a spike extraction *document* for a particular `ndi.element` object, and then, if one is found, plotting the spike waveforms.

**Figure 10.**
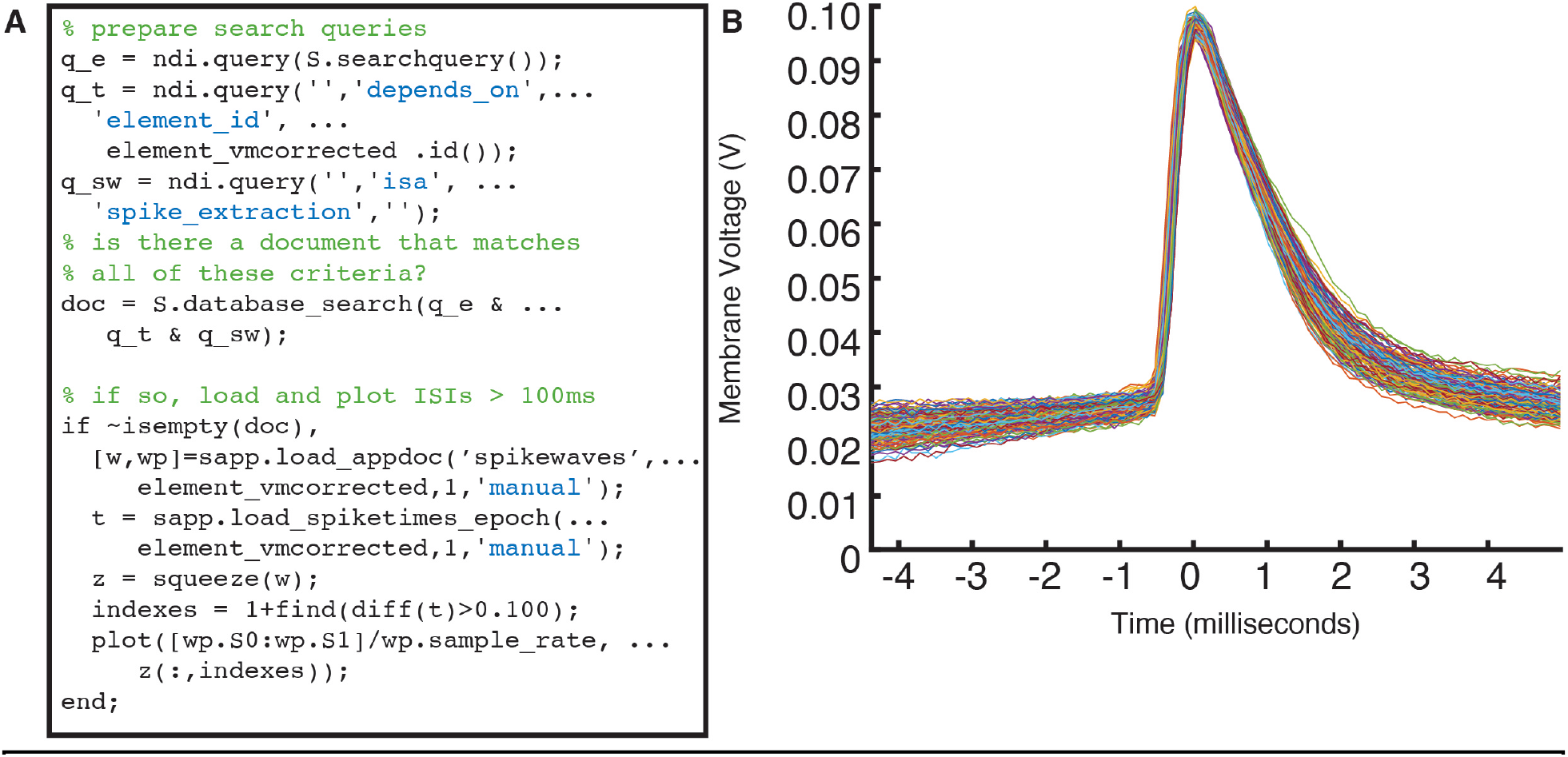
Accessing analysis results involves querying the database with ndi_query. **A)** Code that uses a composition of `ndi.query` objects to look for a document that meets the following criteria: 1) it is of `ndi.document` type ‘spike_extraction’; AND 2) it depends on the `ndi_element` variable named `element_vmcorrected`; and 3) it is from the session `S`. If it finds such a document, then it calls the spike extractor’s method to return the spike waveforms `w` and the parameters `wp`, and spike times `t`. All spikes that have an inter-spike-interval of 100 milliseconds or greater are plotted, as shown in panel **B**.

Developing pipelines in NDI becomes a task of writing small programs that read raw data and/or existing *database documents*, perform computation, and write results back to the *database* in the form of new *documents*. The *documents* exhibit a beautiful structure when plotted as a graph with nodes corresponding to *documents* and edges corresponding to dependencies among *documents*. A representative graph from an experimental session in the study by Roy et al. (2020) is shown in **Figure 11**.

**Figure 11.**
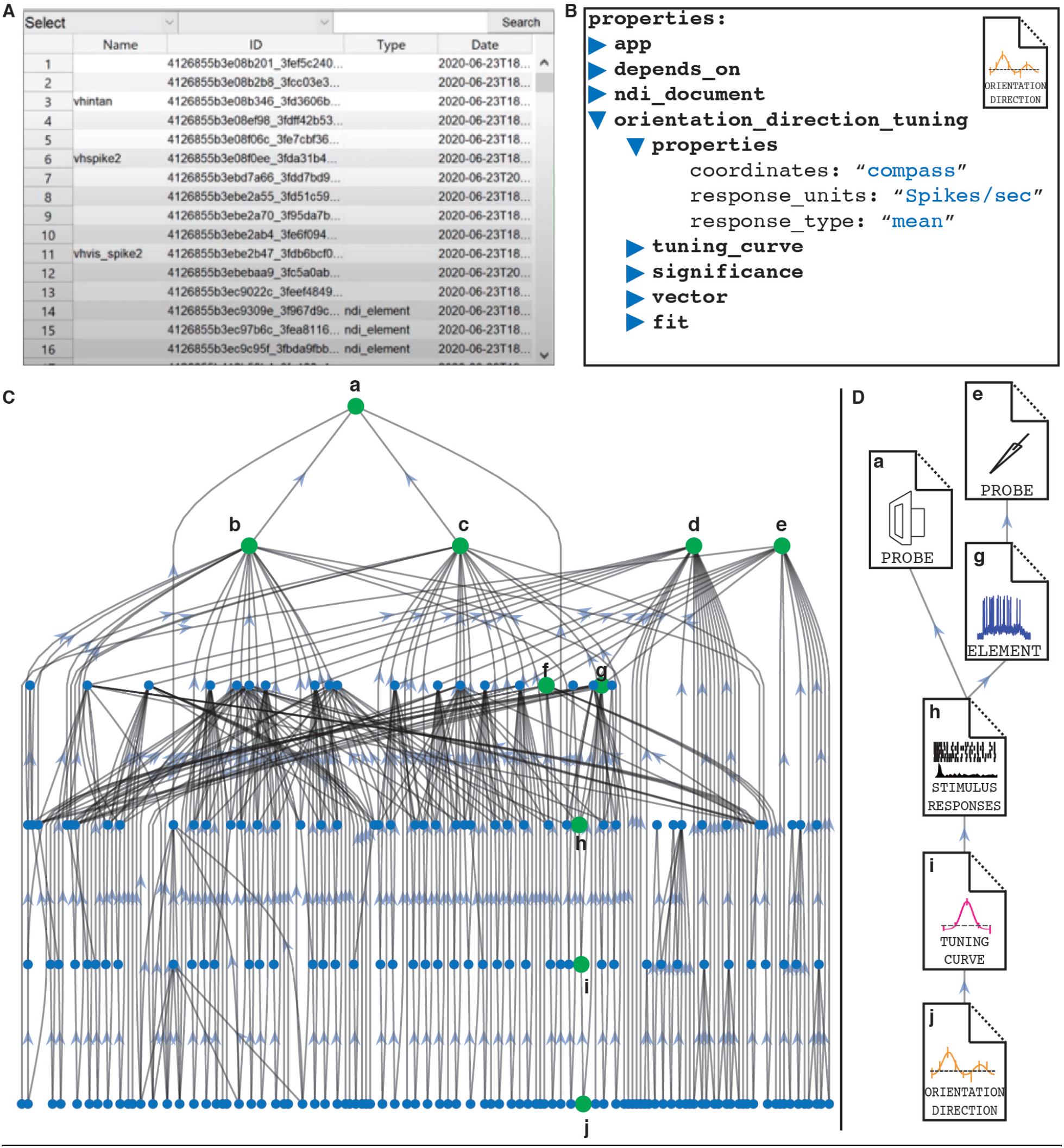
Graph structure of the *database documents* of an example experiment in NDI. **A)** Full graph of documents from an experimental session from Roy et al. (2020). *Documents* are denoted by nodes (blue or green circles), and arrows point from dependent *documents* to the *documents* that they depend upon. In this graph, **a** is a visual stimulus monitor *probe*, and **b** and **c** are stimulus presentation *documents* that describe the presentation of sinusoidal gratings in different directions. **d** and **e** are sharp electrode *probes* corresponding to 2 recordings of different impaled cells. **f** and **g** are *documents* describing the `ndi.element` objects of probe **e** where spikes are removed (**f**) and where spike times are extracted (**g**). **h** is a *document* containing the stimulus responses of the spikes in **g** to the stimulus presentation in **c**. In **i**, these stimulus responses have been collated into a tuning curve. Finally, these responses have been examined to extract orientation and direction index values and to perform a double Gaussian fit, which are all stored in *document* **j. B)** Zoomed in view of the *document* pipeline **a**-**j**.

### Implementation -lower level

The software product implementation of the interface is currently at the level of a working prototype in Matlab and a prototype in Python (see **Materials and Methods**). The low-level database implementation is only a slow prototype, and is currently being modified to use external SQL databases to allow the system to be used at a larger scale. Database documents in the prototype are JSON-based (with a binary blob) but will have stricter typing as the external database options come online. The system has been used to analyze data for a paper (Roy et al., 2020) and will be tested with data from other labs in 2021. The software product is continuously updated on GitHub (see Materials and Methods).

#### Case studies – reading data from many labs

How easy or difficult is it to read data from other labs in NDI? We present in **Figure 12** an example of reading data from 3 laboratories: the Marder Lab at Brandeis (Hamood et al., 2015), the Angelucci Lab at the University of Utah (unpublished data), and the Katz Lab at Brandeis (Mukherjee et al., 2019).

**Figure 12.**
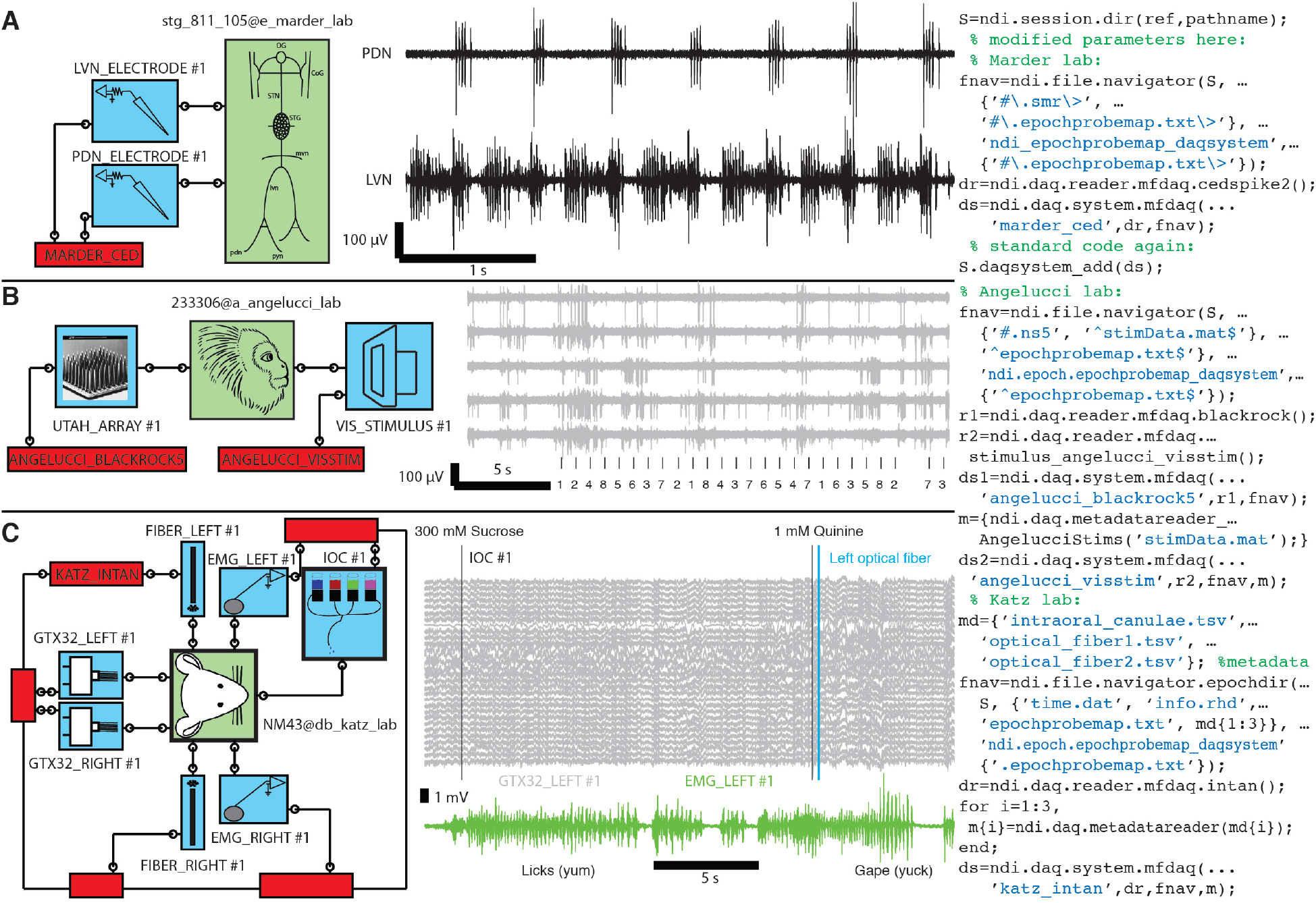
With NDI daq readers and a few parameters, one can read many different types of experiments quickly and directly, without file conversion. *Subjects* (green boxes), *probes* (blue boxes), and *daq systems* (red boxes) are shown. Wires and terminals indicate connections of *probes* to *subjects* and *daq systems*. **A)** Activity of a central pattern generator measured in Eve Marder’s lab (stomatogastric ganglion (STG) of the crab *Cancer borealis*) (Hamood et al., 2015). Electrodes on different nerves indicate the pyloric rhythm that controls the movement of food into the crab’s stomach. The 3 instructions of code needed to specify the *daq system*, modified on a template, are shown at right. Acquisition system was by Cambridge Electronic Design. **B)** Unpublished data snippet from Alessandra Angelluci’s lab showing responses to visual stimulation that were recorded on a 96-channel Utah array implanted in a marmoset. Traces show spikes and numbers, and tick marks are visual stimulus identifier numbers. The 6 instructions needed to set up the 2 daq systems are shown; another 15 lines were needed to build a custom stimulus reader (modified from a similar reader). Acquisition system was by Blackrock Microsystems. **C)** An experiment by Don Katz’s lab (Mukherjee et al., 2019) that explored the relationship between activity in gustatory cortex and whether a rat would choose to consume or expel a taste stimulus delivered through interoral cannulae. The experiment also included optical fibers to optogenetically inhibit neurons projecting to the gustatory cortex from the amygdala. Graph shows EMG recordings (green) indicating licking following sucrose delivery and gaping following quinine delivery. Some inputs to gustatory cortex were inhibited just after quinine was delivered. The 6 instructions needed to express the daq system are at right. Acquisition system was by Intan Technologies. This figure shows how diverse experiments, with different formats and different file organizations, can be read through NDI by specifying only a few parameters. Additional experiments of these types can be read with no new code.

The Marder lab recorded signals from the stomatogastric ganglion of the crab *Cancer borealis*. The lab used a common data acquisition system (Spike2 software from Cambridge Electronic Design), and the data can be specified by creating an `ndi.daq.system` with the `ndi.daq.reader.mfdaq.cedspike2` reader and describing where the files for different epochs are found on disk using an `ndi.file.navigator` object. It requires only 3 instructions (**Figure 12a**) to create the `ndi.daq.system` once, and this `ndi.daq.system` can be used over and over again to access all the data from the experiments in the Hamood et al. study (2015) and many current and past experimental sessions in the Marder lab.

The Angelucci lab recorded 96-channel data from a Utah array in the marmoset (unpublished data courtesy Alessandra Angelucci and Andrew M. Clark). The Angelucci lab used a commercial data acquisition system (from Blackrock Microsystems) and, like many visual labs, use their own visual stimulus system. The Angelucci stimulus system stores its files in Matlab with a time clock that matches the Blackrock Microsystems time clock. For this data, we had to follow a template to make a customized stimulus metadata reader (15 lines of code from a template), and it took 6 instructions to specify the 2 `ndi.daq.system` objects needed to access the Utah array data and visual stimulus parameters and timing data (**Figure 12B**).

The Mukherjee data (2019) included several probes in rat, including dual 32-channel electrode arrays that recorded gustatory cortex bilaterally, dual optical fibers that ontogenetically manipulated activity in gustatory cortex bilaterally, dual EMG electrodes for observing licks and gapes, and intraoral cannulae for delivering tastants directly to the tongue. The Katz lab used a commercial Intan Technologies multifunction data acquisition system, and the code that specifies the `ndi.daq.system` takes just 6 instructions. Again, this `ndi.daq.system` is made once and can be re-used by other members of the Katz lab (**Figure 12C**).

Thus, an analyst who receives data from another lab, regardless of whether that data is packaged in a standard format such as NWB or in custom formats, can gain easy access to the data of other researchers and begin analyses the same day using software that follows the NDI conventions, including apps and custom code. Data that is passed on as an `ndi.session` can be immediately read by other researchers.

## Discussion

We have designed a neuroscience data interface (NDI) that greatly reduces the burden of analyzing datasets from other labs. The interface allows an analyst to quickly address data that is acquired in a variety of formats and stored with a variety of organization schemes on disk. It provides tools for time synchronization across data acquisition systems, and allows experimental *probes* to be addressed directly by the analyst, while the interface performs the necessary reading from underlying *DAQ systems*. The interface contains a *database* that allows experiment objects, analyses, and analyses of analyses to be stored as *documents*, enabling the development of an application ecosystem that performs analysis independently of the format or organization of the underlying data. The results of the dataset can be accessed widely by anyone using the interface, such that the dataset and its analyses are curated for wide distribution.

### An interface with low barriers for curation and exchange

This neuroscience data interface offers several advantages relative to the current neurophysiological data standardization approaches of which we are aware. 1) NDI is grounded in concepts and a vocabulary that is easy for non-coders and coders to grasp. 2) NDI reads data in its native formats, so there are no restrictions for experimental data collection other than a requirement for using a logically consistent scheme and, once, locating or writing an open-source reader for each data type. 3) Reading native formats also offers the significant advantage that the interface can be used regardless of whether the lab performing the data collection wishes or has the expertise to explicitly convert and curate their own data for analysis by others: an experienced data analyst will be able to quickly analyze data using the tools provided by NDI. 4) Reading native formats does not preclude the development of excellent file formats, and implementations of NDI can take partial advantage of fast code created for existing or future formats. 5) There is a *database document* framework so that users and applications can create and abide by *document* templates for saved analyses, so that other users and applications can read and interpret the results of classes of data analyses in a consistent manner. 6) The *database* is scalable and can exist on a user’s computer or in the cloud, and data from multiple experiments can easily be combined in the cloud to form large, searchable *databases* of neuroscience data and analyses. 7) The *database* offers methods for auditing computations and analyses, such that the code and raw data that underlie computations and analyses can be fully tracked and reconstructed. Finally, like many standardization efforts, we aim for the development of an ecosystem of neuroscience analysis apps that will improve reliability, reproducibility, and ease of discovery through re-analysis of data by scientists or amateurs.

### Why not simply a file format?

Why not simply require users to convert their data into a common, standard file format? A standard file format provides several advantages. It provides a common target for development for device manufacturers and for companies and scientists writing analysis software. As the number of channels on some devices become larger, it may be prudent to include hardware in analysis, and a common format facilitates this process. Converting to a common file format also puts the burden of solving the synchronization of different devices outside the scope of the file format, as common file formats such as Neurodata Without Borders (Teeters et al., 2015; Rübel et al., 2019) require the user to import data from various devices into the format, and the scientist performing data analysis is freed from considering these problems.

However, there are many reasons why, in our opinion, a common file format should not be the only tool in our toolbox. The first set of arguments against a common file format is technical in nature. We take it as a given that the most appropriate way to store raw data from an acquisition device (or simulation) will vary according to the particular computational and hardware needs of the device, and these needs may evolve in ways that we cannot imagine at present. For example, the optimal way to compress and store full 3-d voxel images from a calcium imaging experiment involving a major portion of the macaque brain (which may be possible in the future) may be very different from those required to store 3-d voxel images from a 500 µm x 500 µm x 10µm cube. By specifying a common interface standard but leaving the implementation to vary from *DAQ system* to *DAQ system*, we gain most of the benefits of a common file format without the liabilities of imposing a particular storage structure. One may suggest that one could always export the data from a device’s native format to a common file format, but one must remember that a) this is an extra step for the experimenter, and b) this step could be prohibitively expensive (in time) for experiments that require somewhat “online” access to neural responses. Having direct read access via a common reader interface allows the data to be examined “in place” in any file format. Our own experience waiting an hour to convert a few minutes of 1000-channel recordings from a prototype acquisition system in order to perform “online” analysis makes us very enthusiastic about “in place” analysis.

A second set of arguments against a common file format relates to the ease of workflow for the scientists. Our goal was to create a system that can be used at the time of data acquisition. There should be no forced separation between on-line and off-line analysis, so that one can develop best-of-breed tools for either application that do not depend strongly upon the platform or devices being used.

Finally, data curation is clearly a major burden, as there exist file formats that could be used for exchange but very few people use them, although this is improving. The requirement of an extra step at the conclusion of analysis to “export” the data is a barrier to adoption. In NDI, there is no curation step, it is an inherent part of using the data interface.

An interface can bring on board some of the best benefits of an excellent file format, because an interface can read from any file format. As excellent file formats (such as Neurodata Without Borders) are developed, interfaces such as NDI can still read them, and these formats can be used as a target for future development of hardware and software. The NDI approach allows data from these sources to be integrated easily with data from older devices, or newer devices that use a different format for whatever reason (technical, creative, or historical/idiosyncratic). NDI also allows arbitrary time relationships among epochs to be specified and navigated by the interface (local or global), so there are no limits on the data that can be easily included and referenced.

### Stress points: the first DAQ system, ndi.daq.reader, ndi.file.navigator

NDI was designed so that an experienced analyst can specify only a few parameters about the file format (`ndi.daq.reader`) and data organization (`ndi.file.navigator`) in order to get started (**Figure 3**). For most labs, this will entail a small time investment by a user with coding experience to set up the initial *DAQ system* for a lab, or less if the lab uses file formats for which `ndi.daq.reader` objects are already available. After this initial setup, a *DAQ system* definition can be reused as often as necessary, so a majority of lab users will not need this initial expertise.

### Comparisons and synergies with other efforts

This work builds on the experience and expertise of past and current efforts to ease the sharing of data in the neurosciences. A scholarly list of efforts to organize and share neuroscience data is presented in Table 1 of Teeters et al. (2015), and we won’t attempt to enumerate a list of all such projects here. Instead, we will draw comparisons with a few ongoing efforts.

**Table 1:** Key resources table

| Reagent type | Designation | Source or reference | Identifiers | Additional information |
| --- | --- | --- | --- | --- |
| Software | Matlab | The MathWorks, Natick, MA | RRID:SCR_001622 | Software language |
| Software | GitHub | GitHub | RRID:SCR_002630 | Software repository |
| Software | Python3 | www.python.org | RRID:SCR_008394 | Software language |

The idea of an open-source system that can read a variety of file formats is not new. The Matlab project sigTOOL (Lidierth, 2009) and the Python-based projects Neo (Garcia et al., 2014) and SpikeInterface (Buccino et al., 2020) are already capable of reading a wide variety of data formats, and we are using the open source libraries of sigTOOL, Neo, and SpikeInterface extensively in our construction of the Matlab- and Python-based versions of NDI. On top of reading different file formats, NDI adds the ability to deal with different file organizations and explicit management of different timebases on top of managing different file formats or collections. Neo and SpikeInterface manage their raw data output in terms of quantities that are similar to NDI’s epochs.

Neurodata Without Borders (NWB) is an ongoing effort to devise a file format for neuroscience data and analyses (Teeters et al., 2015; Rübel et al., 2019). At present, it requires users to use or write conversion software to save data into a single file that is organized in HDF5 format and that employs a consistent data schema. In NWB, there is no equivalent of the NDI *daq system*; instead, users save what NDI calls *probe* and *element* data directly to the file. The system also offers spaces to save results of “processing” and “analysis”. NWB does not allow for multiple time bases, which simplifies the format greatly for the analyst, but it means that it is difficult to specify situations where probes or other elements have time bases that can be only partially mapped to each other (such as multiple synchronized devices that have only local clocks and no way of mapping to a global time). The format is at present very tied to a file system (1 file per session), although it can be used in conjunction with databases like DataJoint. NWB continues to evolve to broaden its functions and extension capability.

The *document* space of the NDI *database* has commonalities with the tables in the database DataJoint (Yatsenko et al., 2015). For example, the *document* in **Figure 8** can be built by 5 related tables in DataJoint (document classes `ndi_document`,`ndi_epochid`,`ndi_app`,`spikewaves`, `document_class`). Different users may prefer the *table* arrangement of DataJoint or the *documents* of NDI. We designed our *documents* independently of DataJoint and noticed the similarities later. We think that the *document* structure of NDI might be easier for non-programmers to grasp and no more difficult for programmers to query, but the *database* forms share similar forms, including the ability to have dependencies across *table* entries or *documents*. Both DataJoint and NDI lend themselves to the creation of exploration tools that allow users to examine the analyses that have been run and the creation of pipelines – compositions of analyses – that can speed analyses and improve reliability and reproducibility.

### Big challenge: A culture of digital annotation

Although NDI was designed to tackle the heterogeneity of the digital organization of data, our own experience and several colleagues have commented that another barrier to analyzing the data of others is the lack of any consistent digital annotation of data (Teeters et al., 2008; Grewe et al., 2011; Wiener et al., 2016; Sprenger et al., 2019). Often, the only copy of important metadata is written in a physical notebook and is not expressed digitally. Hopefully, as investigators see the utility of common analysis tools, the need to have consistent digital annotations of data and metadata will become clearer and more ingrained in experimental culture.

### Big challenge: Common database schemas for analyses, analyses of analyses

As data interfaces allow more streamlined access to data formats, a new problem arises: how do we read analyses or analyses of analyses from other labs? The database’s flexibility in creating new schemas and *document* types is a double-edged sword. Imagine that one lab develops a set of *database documents* that describes several responses indexes that characterize the response of a neuron to a class of stimuli. Now, imagine that another lab develops its own set of *database documents* for the same purpose, but gives the fields different names and organizes these indexes into a different *document* set. Someone doing a meta-analysis of data from the different labs would either have to recompute the index values from the raw activity of the neurons, or write analysis code that would search the *database* for the *document* schemas of both labs. For example, users are free to design their own schemas in DataJoint, NWB, NDI, or odML (Grewe et al., 2011; Sobolev et al., 2014; Sprenger et al., 2019), but there is no requirement that these schemas be similar or be able to exchange with one another.

Efforts to standardize schemas for certain sub disciplines (such as visual physiologists, or cellular physiologists) could be quite useful, but will take time (Wiener et al., 2016). In our opinions, the development of these schemas have the best chance for broad adoption if they are created independently of software implementation and are not tied to any specific software product. Each software tool may have its own particular advantages for certain applications, and it would be very powerful if users could form queries that make sense across multiple tools. If there were a standard list of metadata for common data types, an interface or file format or database could say it was “ACME 12345”-compliant (where ACME is the name of the organization making the standard, and 12345 was the version of the standard), and users could make common searches across these systems.

The field of fMRI is several years ahead of the physiology and imaging communities in the development of these systems (Cox, 1996; Saad et al., 2006; Gorgolewski et al., 2016; Farber, 2017; Gorgolewski et al., 2017; Nichols et al., 2017; Poldrack and Gorgolewski, 2017).

### Summary

As experimentalists and theorists in neuroscience enter the era of big data, it is necessary to lower barriers of data exchange and to increase access and the ability to search and aggregate data across labs and studies. Some labs have already developed pipelines and tools for exchange of neurophysiology and imaging data (Teeters et al., 2008; Teeters et al., 2015; Yatsenko et al., 2015; Rübel et al., 2019), while the great majority of labs and investigators still use custom or idiosyncratic schemas. Data interfaces allow analysts to quickly work with both types of data, greatly speeding collaborations that might otherwise be too cumbersome. Data interfaces also allow the development of best-of-breed tools that focus on analysis rather than being burdened with the format or organization of the underlying digital data. As more neuroscientists gravitate towards sharing data, utility and ease of use will be important determining factors in adoption and the degree to which users with different levels of computer expertise (users, novice programmers, advanced programmers) can do science with each system. NDI was designed to address all these considerations through conceptual design first, and implementation second, using an interface framework that can reach back into the data of the past and into the data of the future.

## Materials and Methods

### Design of the interface

The neural data interface in its current form was designed and revised over the course of 5 years. The conceptual framework of the system was developed through discussions with Brandeis neuroscience and computer science graduate and undergraduate students. The system began from a Lab Information Management System (LIMS) in the Van Hooser lab, and was rebuilt twice from scratch to incorporate necessary features and simplify the interface and external concepts.

The interface was prototyped in Matlab (The MathWorks) and is available at http://github.com/VH-Lab/NDI-matlab. NDI was used extensively to analyze the data of Roy et al. (2020), and NDI was revised and debugged as necessary to allow a full pipeline analysis. In addition, the process of developing tutorials for user feedback also identified unnecessary complexity and bugs that were revised or simplified. Third party libraries such as sigTOOL (Lidierth, 2009) are extensively used to read a variety of data formats. Functions in NDI also depend on the VH Lab toolbox http://github.com/VH-Lab/vhlab-toolbox-matlab and a set of third-party tools: http://github.com/VH-Lab/vhlab-thirdparty-matlab.

A Python3 version is under construction by SquishyMedia, LLC (Portland, OR). The Python3 distribution is located at http://github.com/VH-Lab/NDI-python. Project neo (http://neuralensemble.org/neo) (Garcia et al., 2014) is used extensively to read a variety of data formats.

## Acknowledgements

This work was funded by an NIH BRAIN Grant (MH114678). We thank members of the Van Hooser lab and the Brandeis systems neuroscience community for comments. We thank Eve Marder’s lab, Alessandra Angelucci’s lab, and Don Katz’s lab for sharing data for demonstration purposes.

